# Morphology of the Ovaries, Uterine Tubes and Uterus of *Pteronotus gymnonotus* (Chiroptera: Mormoopidae)

**DOI:** 10.1101/2022.12.26.521015

**Authors:** Erich Fernando Espinelo Costa, Danielle Barbosa de Morais

## Abstract

The present study aimed to characterize the ovarian, tubal and uterine morphology in the insectivorous bat *Pteronotus gymnonotus*, in specimens collected in the state of Rio Grande do Norte, Brazil. After euthanasia, these organs were histologically processed for inclusion in historesin for morphological and morphometric analysis under light microscopy. The morphological characterization of the ovaries was based on the development of the oocyte and follicle growth, while the uterine tubes were characterized in terms of their anatomy and division of their parts into: infundibulum, ampulla and isthmus, where the height of the tubal epithelium and muscle layer thickness. The analysis of the uterus was based on the formation and thickness of its three layers: endometrium, myometrium and perimetrium. Morphometric analyzes were performed by capturing images of histological slides at different magnifications. The variables were submitted to descriptive analysis, with data expressed as mean and standard deviation. It was observed that the ovaries are bilateral and oval, presenting a squamous to simple cubic epithelium, forming the germinal epithelium, certain stratification regarding the location of the ovarian follicles, where most of the primordial follicles are arranged in the peripheral region of the ovary, however being it is possible to identify many follicles in various stages of maturation in the central region; the uterus is bicornuate and the layers of the uterus and uterine tubes observed follow the same pattern of other eutherian mammals. This information is important to allow comparisons between species, aiming at knowledge about reproductive morphology in mammals, especially those belonging to the order Chiroptera. Therefore, this research is essential to subsidize conservation measures that protect their natural populations, in an effort to maintain the ecological balance.

## 1. INTRODUCTION

Bats are mammals belonging to the order Chiroptera and the only ones capable of true flight. They constitute the second largest order of mammals in number of species in the world, and are present across the globe, with the exception of some remote oceanic islands and polar regions (Nowak, 1999; Fenton and Simmons, 2014).

A great diversity of bats is found in Brazil, with 9 families, 68 genera and 178 species (Nogueira et al., 2014). They inhabit the entire national territory, occurring from the north of the Amazon, through the arid northeast to the mountains of Rio Grande do Sul, and are even present in urban centers. The diversity within this order includes variations in their weight and size where they can be found from bats weighing around 3 grams and having a 15 cm wingspan, such as *Furipterus borrens*, to species that can reach up to 190 grams and a 70 cm wingspan, such as the *Vampyrum spectrum* (Nowak, 1999).

Great diversity is also observed in their eating habits, which can be carnivorous, insectivorous, frugivorous, nectarivorous, polynivorous and hematophagous, and some species may have more than one feeding habit (Fenton, 1992). Thus, fruit bats, for example, are of great biological importance, given that they disperse seeds through their feces and thus contribute to the regeneration of neotropical forests. Insectivores are important controllers of natural insect populations (Boyles et al., 2011).

The great diversity of reproductive patterns found among bats is also notorious, where they have the greatest diversity of reproductive strategies among all orders of mammals (Crichton and Krutzsch, 2000). Females, in particular, show a wide variety in ovarian morphophysiology throughout their reproductive cycle, such as polarized ovaries, preferential ovulation, alternating or double ovulation, including cases of late ovulation; characteristics that may be important in the evaluation of phylogenetic relationships (Crichton and Krutzsch, 2000).

Bats that inhabit temperate regions have only one annual reproductive period, due to the hibernation period that occurs in the winter months. Thus, in the weeks preceding hibernation, follicular growth in the ovaries becomes evident and the production of hormones, such as estrogen and progesterone, becomes significant (Crichton and Krutzsch, 2000). Estrogens will directly influence proliferation, differentiation and follicular development and progesterone will act on the development of the uterus and mammary glands, influencing follicular growth, ovulation and luteinization (Peluso, 2006).

However, bats that inhabit tropical regions do not hibernate and usually no long periods of stoppage in gonadal activity are observed. On the other hand, their reproductive cycles are affected by factors such as temperature, rainfall and food availability (Zortéa, 2007). Thus, not all tropical bats show reproductive seasonality, demonstrating various reproductive patterns, such as seasonal monoestry, characterized by a single reproductive peak per year, bimodal seasonal polyestry, when there are two reproductive peaks during the year, continuous polyestry, with reproductive activity for most of the year, with short periods of inactivity; and aseasonal polyestry, when reproductive activity occurs throughout the year (Willig & Gannon 1993; Zortéa, 2003).

Although most bat populations are in the lowest risk categories, according to the list of endangered species of the International Union for Conservation of Nature and Natural Resources (IUCN, 2019), the increasing destruction of their habitat brings in light of the need to guarantee the maintenance of the stability of bats, as well as their biological control, especially in view of their ecological importance. For this, it is essential to know their reproductive cycles.

However, the scarcity of information about the reproduction of individuals of this order stands out, especially with regard to the morphological and microscopic characteristics of the reproductive organs. When one considers that there are more than 1,000 species of bats described within the order Chiroptera (Simmons, 2005), it becomes evident how limited this knowledge is. An example is the species *Pteronotus gymnonotus*; a small insectivorous bat, whose geographic distribution in South America is restricted (Reis et al., 2007).

Although studies addressing microscopic and physiological aspects are increasing, most studies of the order Chiroptera focus on ecological aspects. Thus, it is necessary to deepen the knowledge about their reproductive biology, in order to generate knowledge about the dynamics of Organs reproductive organs, their peculiarities and reproductive capacity, with a view to maintaining stable populations and species conservation.

## 2. ORDER CHIROPTERA

The Order Chiroptera is represented by the only flying mammals: bats. This order represents the most diverse group of mammals in the world (Nowak, 1999). The word Chiroptera is derived from the Greek *cheir* (hand) and *pteron* (wing), and describes one of the main characteristics of this group, which is having its thoracic limbs adapted into wings. Thus, their hands have five well-developed digits joined by membranes, called patagi; which allow them to truly fly and thus explore the airspace, which consequently allowed them to be widely distributed and dispersed across the planet (Pough, 2008).

This order was commonly divided into the suborders: Megachiroptera and Microchiroptera, the former being a basal, older suborder and the latter the more recent (Simmons and Geisler, 1998). However, recent morphological and genetic investigations have led to a change in this classification. Thus, currently, the order Chiroptera is divided into two clades: Yinpterochiroptera, or Pteropodiformes, and Yangochiroptera, or Vespertilioniformes. The first clade is formed by six families, among them, Pteropodidae and Rhinolophidae (Fenton and Simmons, 2014), while the second is formed by the other families. Among them are Vespertilionidae, Molossidae, Phyllostomidae and Mormoopidae, the first three with the highest number of species found in Brazil (Nogueira et al., 2014).

Bats normally shelter in caves, rock holes, treetops, mines and abandoned buildings, among other places, forming colonies often composed of different species (Reis et al., 2007).

Due to their wide distribution, they stand out for their food diversity, in which they feed on fruits, nectar, parts of flowers, leaves, insects, small fish, amphibians, lizards, birds, small mammals and blood (Fenton, 1992). This diversity of eating habits gives them important ecological roles. Insectivorous bats, for example, are important as insect controllers. It is estimated that some species can eat amounts equivalent to one and a half times their weight in one night (Pavan; Tavares, 2020). Many of these insects are harmful to agriculture and can transmit diseases such as dengue (Nowak, 1999). As a result, insectivores play a fundamental role in the environment, as is the case with the Mormoopidae family, comprising the genera *Mormoops* and *Pteronotus*, which represent good examples of this eating habit (Reis et al., 2007).

Some species of the Mormoopidae family are found only in the Neotropical region, being distributed from southern Mexico to northeastern Brazil. The individuals of this family are characterized by having small eyes, expanded lips and decorated with flaps and folds that form a funnel to the oral cavity when it is open (Pavan; Tavares, 2020). The hair is short and fine, but densely distributed along the body. Its coloration is quite varied, ranging from dark brown to orange. The genus *Pteronotus* comprises seven species: *P. davyi*, *P. gymnonotus*, *P. macleayii*, *P. parnellii*, *P. personatus*, *P. pristinus*, *P. quadridens*. The species *P. davyi* and *P. gymnonotus* differ from the rest of the genus in that their wing membrane is attached to the body at the line of the vertebral column, covering the fur on the back, giving the impression that it has no hair (Reis et al., 2011).

*Pteronotus gymnonotus* can be found from Mexico, reaching South America, through Peru, Bolivia, Guianas and Brazil (Figure 1). Males and females have a similar average head and body length, around 64 mm, and an average weight of approximately 12.6 g for males and 13.6 g for females. A feature of this species is its refuge in caves, where it forms colonies with thousands of individuals, along with other species of the same family. Their diet is basically based on insects, mainly beetles, flies and moths (Lopez-Wilchis *et al*., 2021).

**Figure 1.**
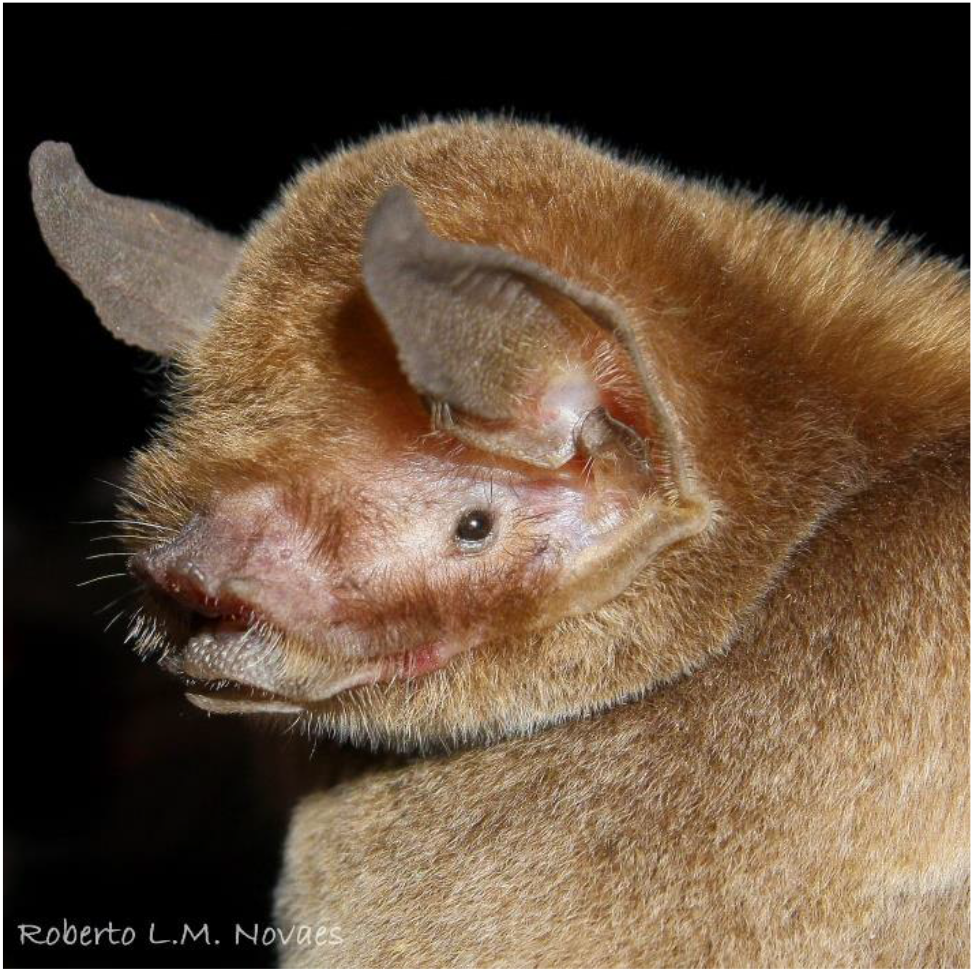
Specimen of *Pteronotus gymnonotus* belonging to the Mormoopidae family. Photo: Roberto Novaes.

### 2.1 REPRODUCTION IN BATS

The reproductive system of bats basically follows the same pattern as that of most eutherian mammals. Females have a pair of ovaries and fallopian tubes, connected to the uterus and vaginal canal (Pough, 2008).

The ovaries are generally polarized, where their surface epithelium is restricted to the medial side of the ovary and primordial-type ovarian follicles are limited to an immediately adjacent zone, taking the example of the species *Glossophaga soricina* (Komar et al., 2007).). Ovulation can be alternating or double (Griffin et al., 2006). During the development of ovarian follicles, the stages are observed: primordial, where the oocyte is surrounded by a single layer of squamous follicular cells; primary unilaminar, in which the oocyte is surrounded by more than one layer of cubic granulosa cells and the theca begins to develop; primary multilaminar, when the oocyte is surrounded by more than one layer of granulosa cells; secondary, when the granulosa layers surrounding the oocyte present an antrum; and mature, where the antrum is wide and the oocyte is eccentric, covered by the *corona radiata* and supported by the *cumulus oophorus* (Griffin et al., 2006). The latter becomes dominant and goes through the ovulation process, where there is the release of the oocyte that is ready to be fertilized.

The ovulation process is formed by a series of morphological and biochemical events within the mature follicle. Therefore, several genes are activated in the ovaries, leading to enzymatic and structural changes under the influence of gonadotropins and sex steroids that are modulated by various growth factors. All these events ensure that the oocyte is expelled from the ovarian surface, and the remaining structures of the mature follicle form the corpus luteum (Beguelini et al., 2020). Thus, after the rupture of the follicular wall, tissue reorganization occurs, causing the differentiation of granulosa cells and theca cells into luteal cells, originating the corpus luteum (Rodrigues, 2019).

This gland is composed of functional cells for the synthesis of progesterone, the main regulator of the pituitary secretion of gonadotropins necessary for the maintenance of the corpus luteum until the initial pregnancy (Bueno, 2019). If it does not occur, the corpus luteum regresses and begins its degeneration process, and is now called the hemorrhagic corpus luteum (Bueno, 2019). As its regression advances, a scar tissue called corpus *albicans begins to form* (Beguelini et al., 2013), culminating in a decrease in the production of estrogen and progesterone by the corpus luteum, which contributes to the elimination of the functional layer. of the endometrium. It is noteworthy that in most species reabsorption of this tissue occurs; however, in some species, such as *Carollia perspicillata* and Glossophaga soricina, true menstruation may occur, thus initiating a new cycle (Rasweiler; Badwaik; Mechineni, 2010).

The newly ovulated oocyte will be captured by the uterine tubes, which are bilateral, attached to the uterus or uterine horns. They are generally short and tortuous (Reis, 2011), but have the same pattern as other eutherian mammals, including the division of their parts, which are divided into the infundibulum, located at the ends, adjacent to the ovary and opens into the peritoneal cavity near the ovary. ovary with extensions called fimbriae; ampulla, portion located just after the infundibulum, a region with a thinner wall and numerous folds; isthmus, medial portion of the tube, generally of smaller caliber, its muscular wall is thicker and also has fewer and less branched folds than the ampulla region; and an intramural part, located at the other end that crosses the uterus and opens inside the same organ. (Rasweiler et al., 2011). The main function of the uterine tubes is to transport female and male gametes and ensure an adequate and conducive environment for fertilization and the initial development of the zygote (Reis, 2011).

They are composed of four basic layers: the serous layer, formed by connective tissue and mesothelium; a muscle layer formed of smooth muscle arranged in an inner circular layer and an outer longitudinal layer; and the mucosa formed by the epithelium and a lamina propria of loose connective tissue (Beguelini et al., 2020). The epithelium is usually simple and varies from cubic to columnar (Beguelini et al., 2013), where it is composed of two cell types, secretory and ciliated cells, but in the vast majority, secretory cells are rare, and the epithelium is formed mainly, by hair cells that move towards the uterus, taking in that direction the mucus that covers its surface, produced by the secretory cells that are interposed between the hair cells (Bueno, 2019; Beguelini et al., 2020).

At the time of ovulation, the infundibulum tapers and approaches the surface of the ovary, favoring the capture of the ovulated oocyte, providing the fimbriae to facilitate the capture of the oocyte after it leaves the ovary. Generally, the meeting of the sperm with the oocyte occurs at the isthmus-ampulla junction, where fertilization occurs. The zygote is then transported to one of the uterine horns or uterus, where implantation will occur (Bueno, 2019).

The uterine morphology of bats is variable, and can be double, bipartite, bicornuate or simple (Rasweiler and Badwaik, 2001; Komar et al., 2007). Some species from tropical regions present true menstruation, such as the fruit bat *Rousettus leschenaultii* belonging to the family Pteropodidae (Zhang et al., 2007) and also in species from temperate regions such as *Myotis ricketti*, belonging to the family Verpertilionidae (Wang et al., 2008). Gestation can vary between species, but on average lasts 44 days. Some females are capable of giving birth to up to five offspring per gestation, although this is a rare finding (Reis et al., 2007).

While studies on the morphology of the reproductive system are scarce, there are numerous studies on the social organization and reproductive patterns, which are based mainly on observing the periods of the year when there are pregnant and/or lactating females. Thus, it is known that bats from temperate regions have seasonal reproduction and are monoestric, since they face periods of hibernation related to low temperatures in winter (Barros, 2013).

The reproduction of bats in the tropical region, although they do not hibernate, is influenced by abiotic factors, such as rainfall, temperature and food availability, among others (Carvalho et al., 2019). For this reason, these bats can be monoestrous, polyestrous and even reproduce continuously throughout the year (Zortéa, 2003). This diversity of reproductive strategies, combined with its dietary diversity and ability to perform true flight are joint factors that contribute to the great evolutionary success of this order, which is evidenced by the great richness of existing species and its vast area of occupation on the globe. (Lopez-Wilchis *et al*., 2021).

The great reproductive diversity found among bats is instigating, stimulating research regarding morphological aspects, given the scarcity of existing studies. The lack of studies on the reproduction of *Pteronotus gymnonotus is highlighted*, as well as on any body system of the species. This information is important to allow comparisons between species, including other mammals, aiming at knowledge about reproductive morphology in mammals. And considering the order Chiroptera, this knowledge is fundamental to subsidize conservationist measures that protect their natural populations, in an effort to maintain the ecological balance.

## 3. MATERIALS AND METHODS

### 3.1 STUDY AREA AND ANIMAL COLLECTION

The captures were carried out in the municipality of Felipe Guerra, Rio Grande do Norte, Brazil, after obtaining a license, No. 62269-1, from the Chico Mendes Institute for Biodiversity Conservation (ICMBio). Only adult females were captured.

After obtaining the license, No. 017/2018, from the Ethics Committee for the Use of Animals of the UFRN (CEUA), euthanasia was performed in the field, shortly after the capture of the animals. Thus, after weighing on a pesola-type scale, the animals were euthanized with 240 mg/Kg of Ketamine and 150 mg/Kg of Xylazine, intraperitoneally, and the physiological parameters were observed until the cessation of heartbeats and respiratory movements. After death was confirmed, the reproductive organs were collected. This research was also registered in the National System for the Management of Genetic Heritage and Associated Traditional Knowledge, No. A9CEAd0 (SisGen).

adult females of *Pteronotus gymnonotus were collected*, identified based on the observation of the fusion of the epiphyseal cartilage of the fourth finger, at the metacarpal-phalangeal junction (Kunz and Anthony, 1982). The reproductive status of each female was assessed at the time of capture, and those with obvious pregnancy and/or lactation characteristics (evident mammary glands and nipples and abdominal inspection/examination) were separated and excluded (Zortéa, 2003).

### 3.2 HISTOLOGICAL PROCESSING

The reproductive organs were fixed in a 4% Paraformaldehyde solution for 24 hours and stored in 70% Ethanol. The ovaries, uterine tubes and uterus were histologically processed for inclusion in Glycol-Methacrylate (Historesina®, Leica). Sections with 3 μm thickness were performed in an automatic rotary microtome, in semi-serial cuts to obtain the best visualization of the structures (Leica RM2255), and the slides were stained with Hematoxylin-Eosin to carry out the morphological and morphometric analyses.

### 3.3 MORPHOLOGICAL AND MORPHOMETRIC ANALYSIS

To carry out the morphometric analyses, images of the histological slides were captured at different magnifications, using a photomicroscope (BEL Bio2/3 Eurekam 5.0). For measurements, Image-Pro Plus software, version 6.0 for Windows, was used, where all measurements were performed individually for each ovary. Assuming the spherical shape of the ovaries of P. gymnonotus, the diameters were calculated using the following equation:

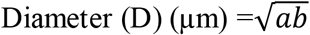

Where (a) refers to the measurement of the diameter of the ovaries and (b) the diameter at a 90° angle (Willians, 1977). The mean diameter (D) was calculated for each ovary and later used to calculate the volume:

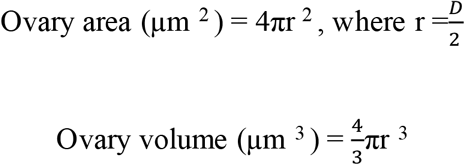

The frequency of follicle types was evaluated in at least five cuts for each animal, starting with the first cut and the others at every multiple of five, that is, the 1st, 5th, 10th and so on. The frequency was estimated by counting follicles with the oocyte nucleus included in the section plane, and the relative percentage was estimated by calculating the proportion of each follicle, expressed as a percentage of the total population.

To estimate the volumetric proportion of the ovarian components, the same Image-Pro Plus software, version 6.0, was used, according to the procedure by Weibel et al. (1978), using the point grid system with 130 points in total for each animal. The relative percentage (%) was calculated after counting the number of points that coincided with each ovarian component. The characterization of the ovarian follicles was performed, based on oocyte development and follicle growth, basically in five stages: primordial follicles, unilaminar primary follicles, multilaminar primary follicles, secondary follicles and mature follicles, in addition to the corpus luteum (Van Den Hurk, 2005). In addition, morphometric analyzes were performed, starting with the absolute frequency of follicles, diameter of follicles and oocytes, and estimation of the number of follicles per ovary (Bueno et al., 2019). The diameter of the follicles was measured based on the surface of the first layer of follicular cells passing through the central region of the oocyte, taking into account the region where the cell nucleus is located. For measurements, Image-Pro Plus software, version 6.0 for Windows, was used, where all measurements were performed individually for each follicle and oocyte. The same formula for the diameter of the ovary was used, assuming the spherical shape of the follicles.

The uterine tubes were analyzed, based on the division of their parts, into three distinct regions: infundibulum, ampulla and isthmus. In addition, the thickness of the muscle layer and the height of the epithelium were analyzed (Beguelini, 2020). The thickness of the muscle layer and height of the epithelium were evaluated using Image-Pro Plus Software, version 6.0. The analysis was performed in five cross-sections, taking as reference the lumen of each region (infundibulum, ampulla and isthmus), per animal, obtaining the final value as the average of 6 measurements in a concurrent straight line in different positions of each part of the tube. Based on the muscular layer, the distance between the free surface of the mesothelium of the serosa and the limit of the lamina propria, where the connective tissue appears; and for the height of the epithelium, the distance between the surface facing the lumen and the edge of the adjacent lamina propria.

The uterine morphology was analyzed, based on the formation and thickness of its three layers: externally, the perimetrium, consisting of mesothelium and connective tissue, the myometrium, consisting of smooth muscle, and the endometrium, the uterine mucous layer (Beguelini, 2020). Thus, the thickness of the endometrium and myometrium was analyzed, based on the endometrium, the distance between the free surface of the endometrial epithelium and the limit of the lamina propria with the myometrium; and for the myometrium, two layers of smooth muscle tissue, inner circular and outer longitudinal, were considered.

The thickness of the uterine layers (endometrium and myometrium) were evaluated using Image-Pro Plus Software, version 6.0. The analysis was carried out in ten cross-sections, close to the center of the organ, per animal, obtaining the final value as the average of 10 measurements in a parallel and concurrent straight line in different positions of each layer of the uterus.

The results obtained were submitted to descriptive analysis, and the values obtained regarding the morphometric analysis of the ovaries, uterine tubes and uterus expressed by means and standard deviations.

## 4. RESULTS

### 4.1 GENERAL MORPHOLOGY AND MORPHOMETRY OF THE OVARIES

The ovaries of *Pteronotus gymnonotus* are bilateral oval structures located in the pelvic region, supported by ligaments and communicating with the uterus through the uterine tubes.

Histologically, the ovaries of *P. gymnonotus* were made up of two distinct regions: the central region of the ovary, formed by loose connective tissue containing some blood vessels and lymphatic vessels; and the peripheral region, containing most of the ovarian follicles in various stages of maturation inserted in connective tissue, in addition to the corpus luteum. The limit between the central region and the peripheral region did not present a clear delimitation.

The surface of the ovary was covered by an epithelium that varied from squamous to simple cubic, forming the germinal epithelium. Underlying this, a layer of dense connective tissue, called tunica albuginea, was observed between the ovarian epithelium and the underlying cortex (Figure 2).

**Figure 2.**
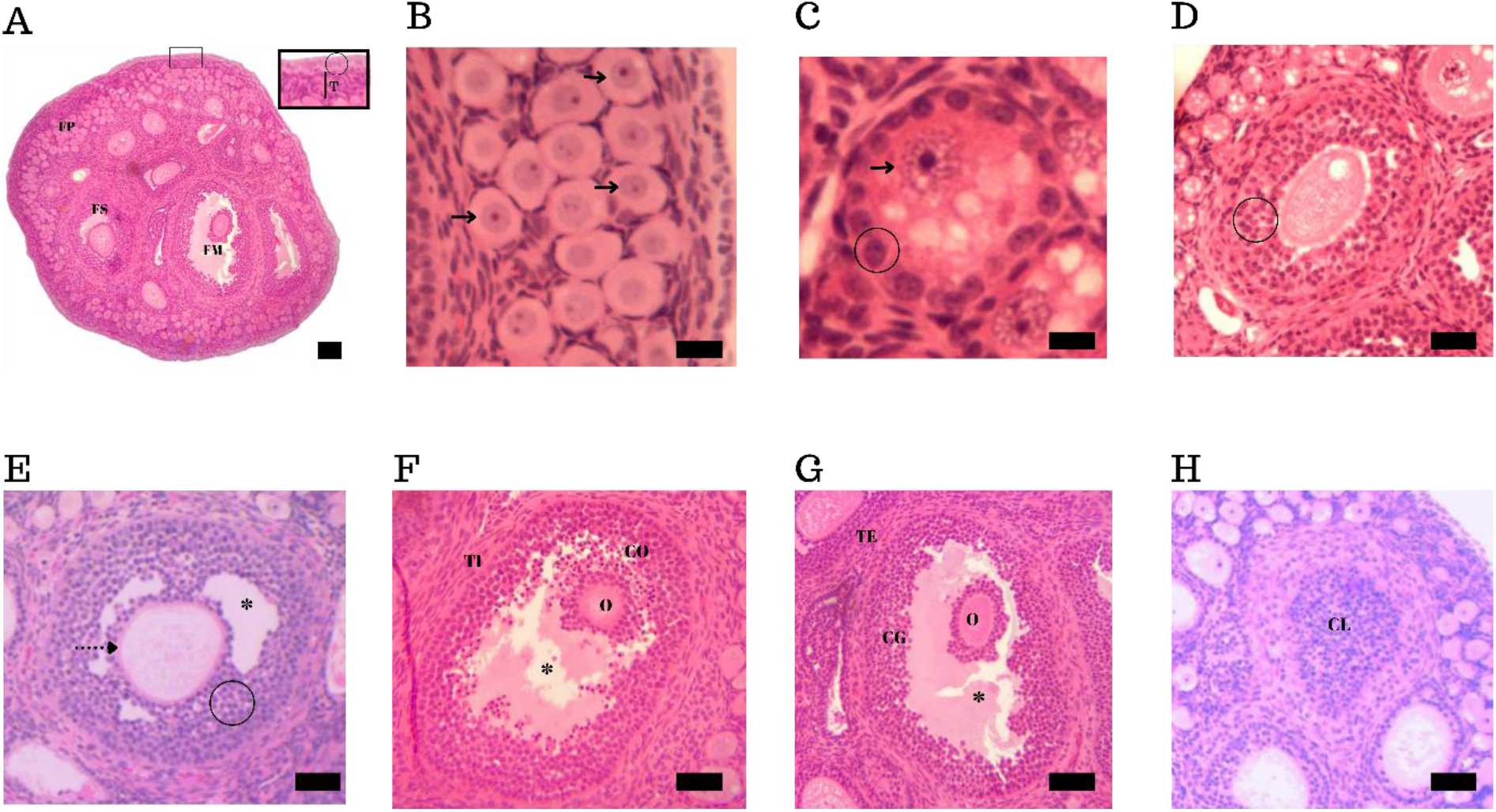
Cross-sections of the ovaries of *Pteronotus gymnonotus*. A: Ovary with primordial follicle (FP), secondary follicle (FS) and mature follicle (FM), in addition to a section of the epithelium, where we have the tunica albuginea (T); B: Primordial follicle with the oocyte (black arrow); C: Unilaminar primary follicle with only one layer of follicular cells (circle); D: Multilaminar primary follicle with several layers of follicular cells (circle); E: Secondary follicle with the formation of the antrum (*) and the formation of the zona pellucida (dotted arrow); F: Mature follicle with its oocyte (O) adhered to *Cumulus Oophorus* (CO); and the Internal Teca (IT); G: Mature follicle with the antrum (*) and its granulosa cells (CG) and the Theca Externa (TE); H: Corpus Luteum. Staining: Hematoxylin-Eosin (HE). Bar: 18 μm.

A certain stratification was observed regarding the location of the ovarian follicles, where most of the primordial follicles were arranged in the peripheral region of the ovary, although it was possible to identify many follicles in various stages of maturation in the central region (Figure 2).

Primordial follicles were characterized by having a primary oocyte surrounded by a single layer of flattened follicular cells, as can be seen in figure 2 – A and B. Most were found in the peripheral region, however a part was also located in the central region.

With its development, these surrounding flattened cells proliferate and become cubic, but still remain arranged in a single layer and the increase in size of the oocyte can be observed. At this moment, the follicle is called unilaminar primary follicle, as it only has a single layer of cuboidal follicular cells (Figure 2 – C).

After successive mitotic divisions, the single layer of follicular cells forms a stratified epithelium, with multiple layers of granulosa cells, characterizing a multilaminar primary follicle, surrounded by the zona pellucida. It is at this stage that a thick amorphous layer, called the zona pellucida, is secreted and surrounds the entire oocyte (Figure 2 – D).

With the development of follicles, mainly due to the increase in size and number of granulosa cells, these grow and occupy deeper areas of the cortical region. At this moment, the follicular fluid begins to accumulate between the follicular cells and small cavities begin to appear, which, when united, will form the antrum (Figure 2 – E). At this stage, the follicle is called a secondary follicle.

During this reorganization of the granulosa cells, some cells of this layer are concentrated in a certain place on the follicle wall, forming a small thickening, the *cumulus oophorus*, which serves as a support for the oocyte. In addition, a small group of follicular cells surround the oocyte, constituting the *corona radiata*. It is at this moment, when one follicle grows much more than the others, that the dominant follicle is formed, which can reach its maximum stage of development and continue until ovulation, therefore being called a mature follicle (Figure 2 – F and g).

The corpus luteum (Figure 2 - H) was observed in most of the ovaries, indicating that the ovaries are functional, that is, they are able to ovulate, as can be seen in Table 2, giving an average percentage of approximately 10% of the area of the ovaries. ovarian components. Only the corpus luteum was observed at the beginning of its formation, where it was possible to verify the granulosa cells invading the entire antrum and being compacted by the internal and external theca, thus initiating the process of changes to luteal cells and initiating hormone synthesis.

**Table 1.** Mean diameter of *Pteronotus gymnonotus ovarian follicles*. Data expressed as mean ± standard deviation.

| | primordial follicle<br>($\mu\text{m}$) | | unilaminar primary follicle<br>($\mu\text{m}$) | | multilaminar primary follicle<br>($\mu\text{m}$) | | secondary follicle<br>($\mu\text{m}$) | | mature follicle<br>($\mu\text{m}$) | |
| --- | --- | --- | --- | --- | --- | --- | --- | --- | --- | --- |
|  | DF | OF | DF | OF | DF | OF | DF | OF | DF | OF |
| Average | 26.70 | 11.91 | 65.94 | 14.74 | 123.29 | 50.11 | 156.35 | 64.03 | 270.31 | 74.41 |
| DP | 2.00 | 1.70 | 11.19 | 9.23 | 31.33 | 31.28 | 35.76 | 26.32 | 47.13 | 1.62 |

DF = Follicle diameter. DO = Oocyte diameter.

**Table 2.** Ovary components, in percentage, of *Pteronotus gymnonotus*. Data expressed as mean ± standard deviation.

|  | primordial<br>follicle<br>BR | unilaminar<br>primary<br>follicle<br>BR | multilaminar<br>primary<br>follicle<br>BR | secondary<br>follicle<br>BR | mature<br>follicle<br>BR | tunica<br>albuginea<br>BR | stroma<br>BR | corpus<br>luteum<br>BR |
| --- | --- | --- | --- | --- | --- | --- | --- | --- |
| Average | 19.24 | 10.06 | 22.07 | 10.76 | 14.62 | 9.14 | 17.60 | 9.39 |
| DP | 7.38 | 6.57 | 8.30 | 9.19 | 6.17 | 1.64 | 6.61 | 2.35 |

Table 1 presents the mean values referring to ovarian morphometry. It was possible to observe that the average diameter of the oocyte in the primordial follicle represented almost half of the total diameter of the same follicle, which was not repeated in the diameter of the other follicles. It was visible that the average diameter of the follicles increased as there was growth and development, as can be seen in the average diameter of secondary and mature follicles (156.35 μm and 270.31 μm, respectively). The average oocyte diameter also increased with the general development of the follicles, although to a lesser extent, as the average oocyte diameter in primary multilaminar follicles and in the mature follicle (means of 50.11 μm and 74.41 μm, respectively).

Table 2 presents the mean values referring to the ovarian components, showing the mean percentage of each one. It was possible to observe a predominance of multilaminar primary follicles, with an average of 22%, followed by primordial follicles with an average of 19%. Thus, together, these two types of follicles represented about 41% of the ovarian components in the specimens used in this study. The corpus luteum had the lowest percentage among the ovarian functional structures, with an average of 9.39%. Secondary and mature follicles together accounted for an average of 25.38% of the ovarian area.

In the comparison between the ovaries with and without the presence of the corpus luteum, it is noticed that the diameter, area and volume of the ovary without the corpus luteum are smaller than in the ovary with the corpus luteum, showing, once again, the process of regression of the corpus luteum. ovary that did not ovulate (Figure 3).

**Figure 3.**
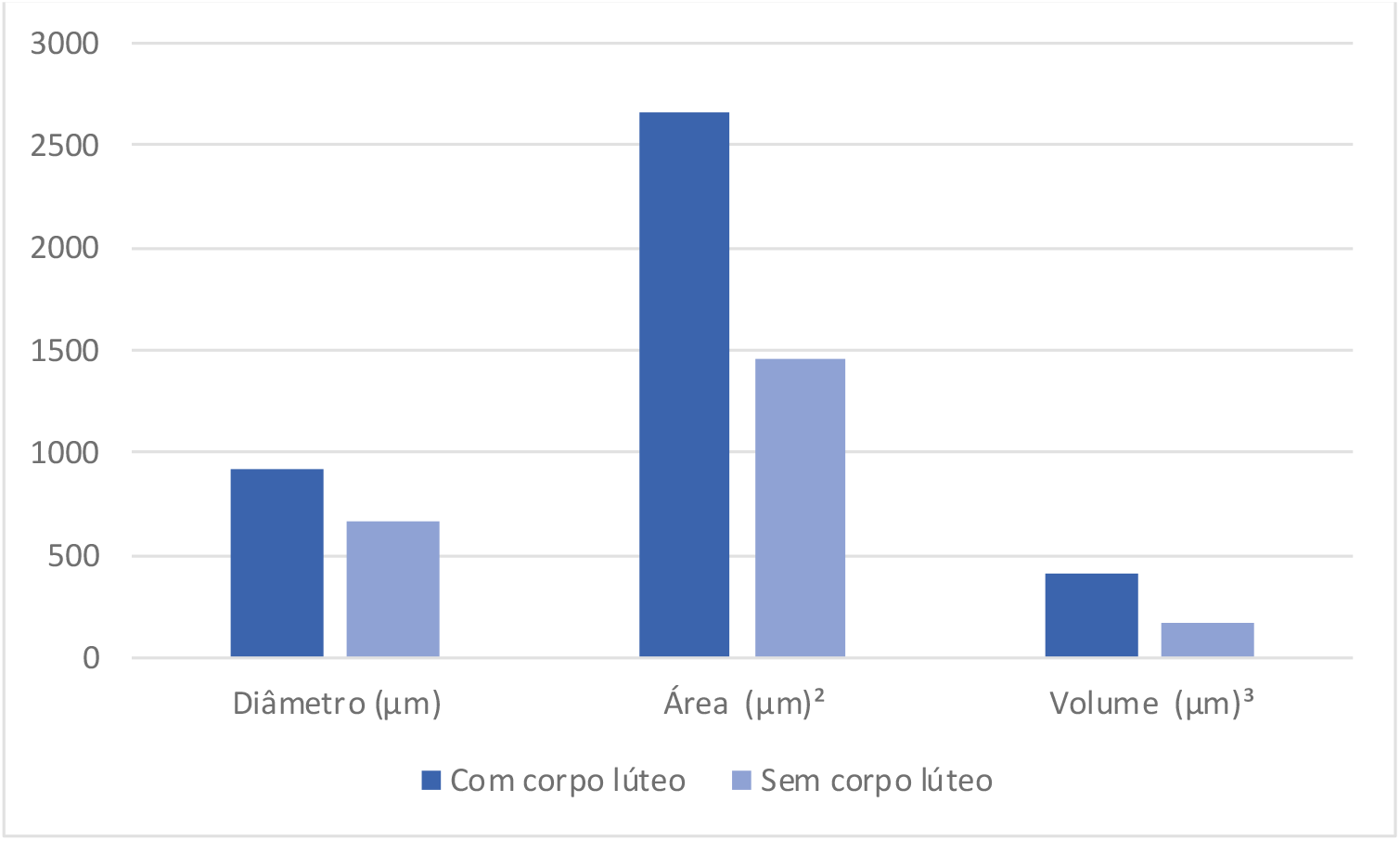
Comparison of the diameter, area and volume of the ovaries of *Pteronotus gymnonotus*, with and without the presence of a corpus luteum.

### 4.3 GENERAL MORPHOLOGY AND MORPHOMETRY OF THE FALLOPUS TUBES

The uterine tubes were short and tortuous, with a funnel-shaped expansion, the infundibulum. This closes into an ostium, the abdominal ostium of the fallopian tube, which is continuous with a more dilated portion of the tube, the ampulla. This goes towards the uterus, with the isthmus, becoming narrower until its junction with the uterus (Figure 4), and the intramural part, where the tube connects with the uterus, which was not observed in the animals analyzed in this study.

**Figure 4.**
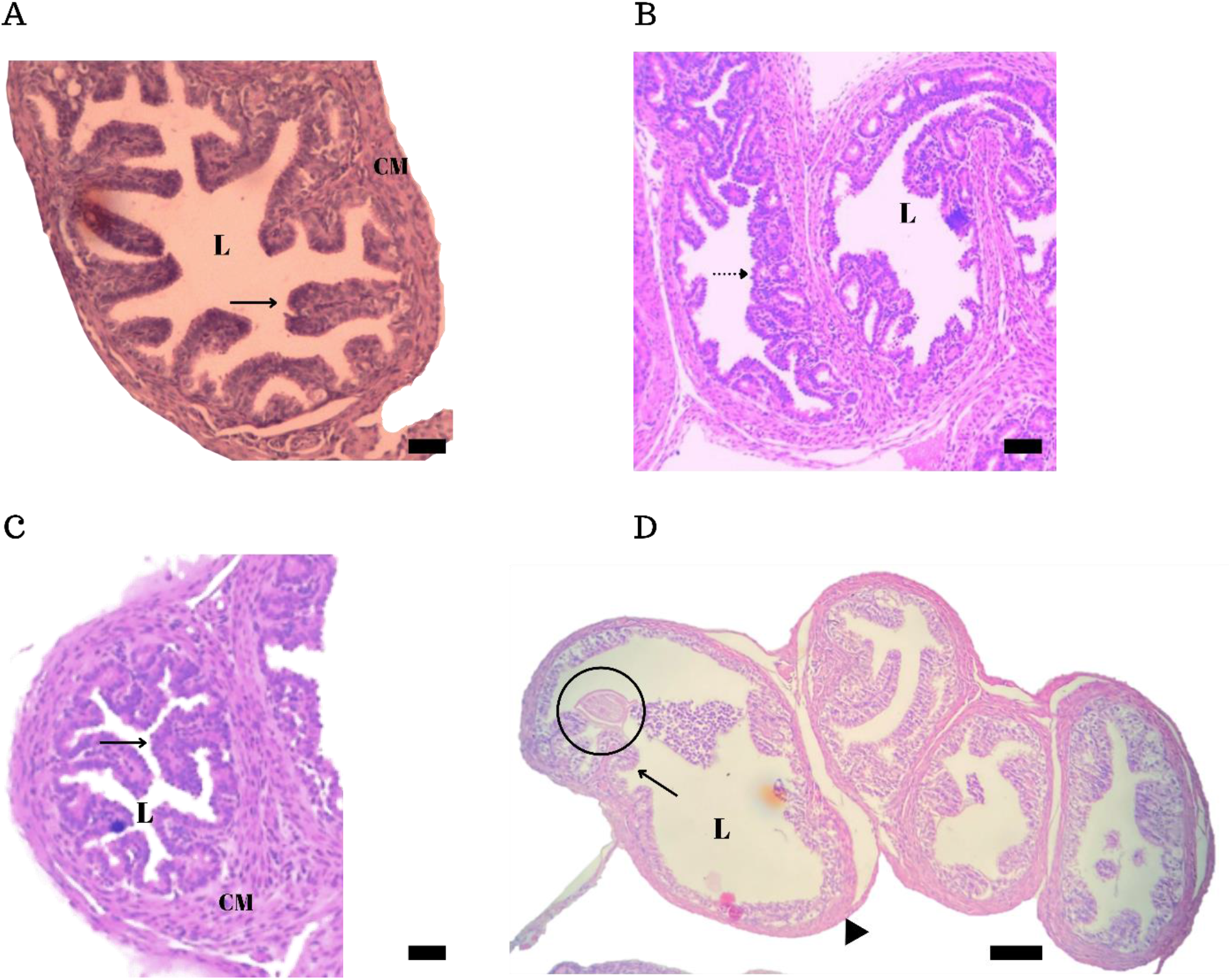
Fallopian tube of *Pteronotus gymnonotus*. A: Infundibulum with visible lumen (L), its folds (black arrow) and with the muscular layer (CM); B: Ampoule with visible lumen and its longitudinal mucosal folds (dotted arrow) showing cilia; C: Isthmus showing its folds (black arrow) and muscle layer (CM); D: Cross section of the uterine tube with visible lumen and an oocyte adhered to the mucosal folds (circle); and the serous layer (arrow head). Staining: Hematoxylin-Eosin (HE). Bar: 60 μm.

Histology of the uterine tube showed three specific layers: the serosa; a muscular layer; and a mucous layer that varied according to the parts of the tube. The serosa being formed of connective tissue and mesothelium; the muscle layer formed of smooth muscle and arranged longitudinally; and the mucosa showed a lot of folding in all its parts, but mainly in the isthmus, with less visibility of the lumen being verified; in addition, the mucosa presented epithelium and a lamina propria of loose connective tissue. The epithelium was simple in the analyzed animals, ranging from cubic to columnar, with interspersed ciliated and non-ciliated cells being observed.

Table 3 presents the mean values referring to the morphometry of the fallopian tubes of *P. gymnonotus*. The thickness of the muscular layer between isthmus, ampulla and infundibulum did not undergo considerable variation in the specimens studied, so that the average between them was 21 to 25 μm. However, variations in the height of the epithelium and in the appearance of the mucosa were observed, thus making it possible to distinguish between its anatomical portions.

**Table 3.**
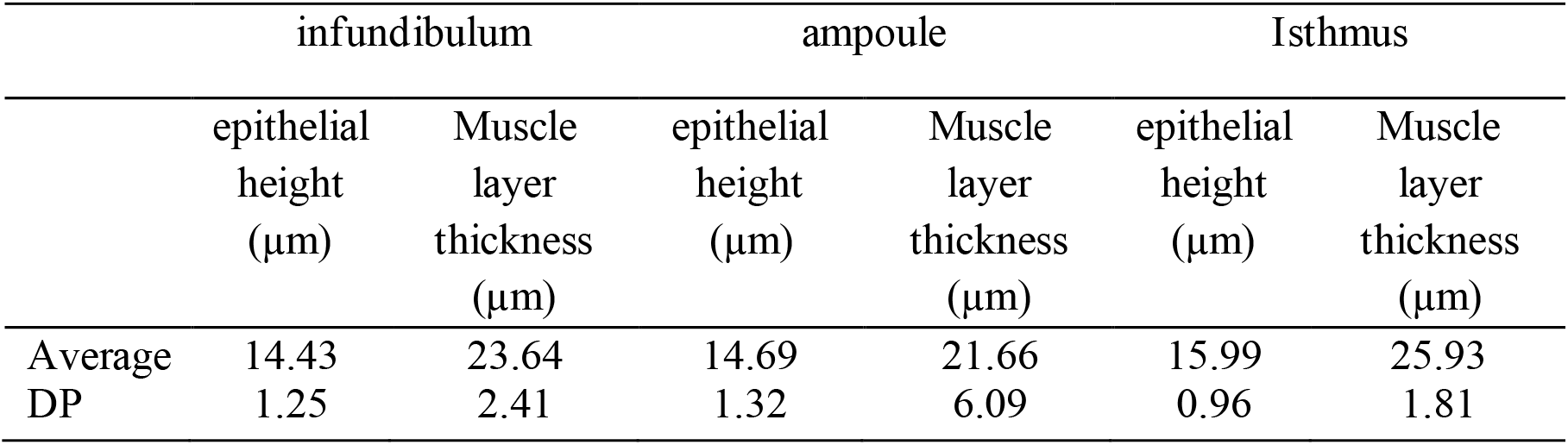
Components of the Uterine Tube of the bat *Pteronotus gymnonotus*. Data expressed as mean ± standard deviation.

### 4.4 GENERAL MORPHOLOGY AND MORPHOMETRY OF THE UTERUS

*P. gymnonotus* females have two lateral uterine processes, thus characterizing a bicornuate uterus; which unite in the caudal portion, forming a single uterine body. This ends in a denser portion of the wall, the uterine cervix. The ovaries were connected to the uterus through the uterine tubes, in the cranioventral portion of each uterine horn (Figure 5 - A).

**Figure 5.**
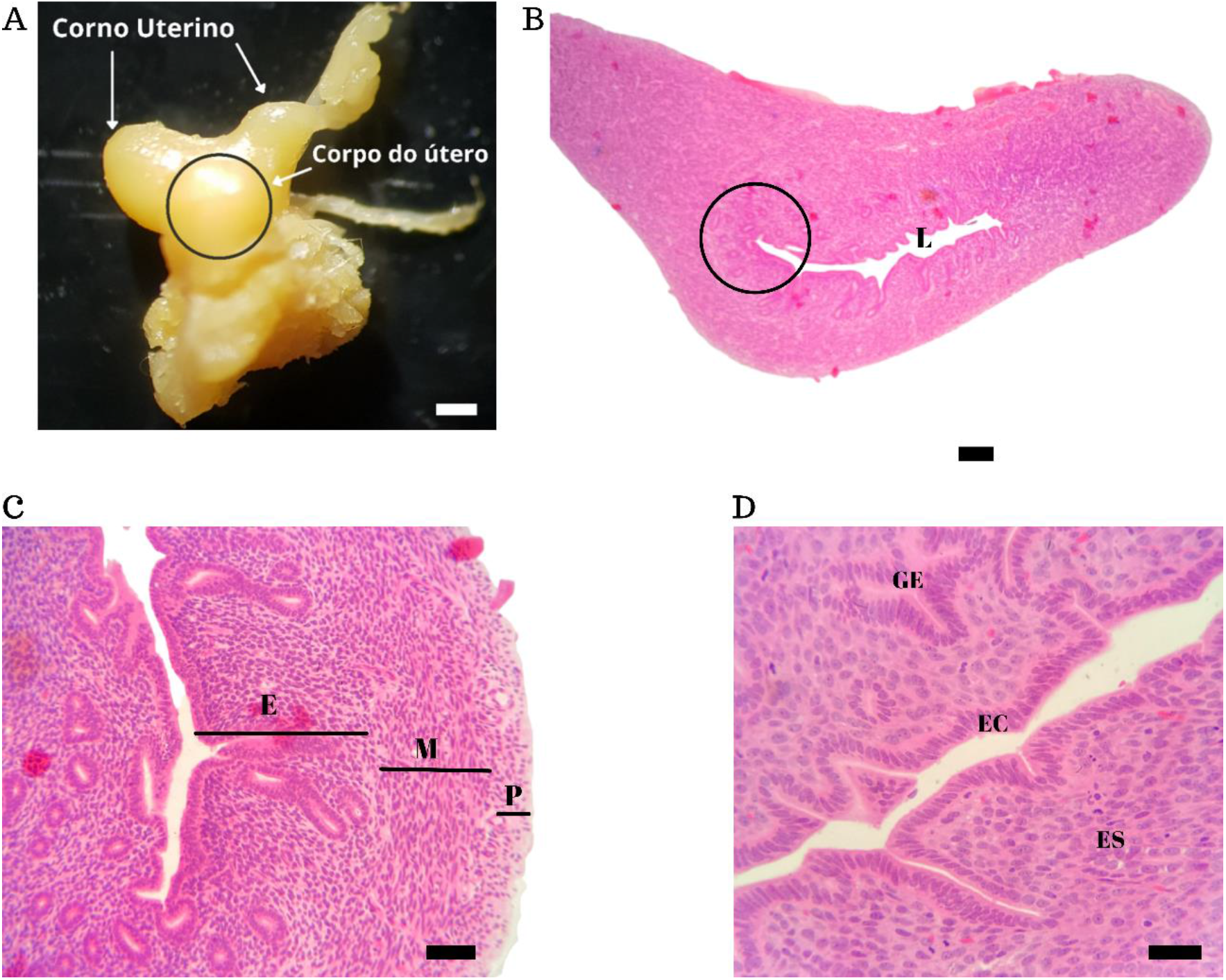
Uterus of *Pteronotus gymnonotus*. A: Anatomy of the uterus with the uterine horns (White bar = 7 mm). B: Bicornuate uterus, with visible lumen. C: Subdivisions of the uterine wall into endometrium (E), myometrium (M) and perimetrium (P). D: Endometrium with Simple Columnar Epithelium (EC), endometrial glands (GE), stroma (ES). Staining: Hematoxylin-Eosin (HE). Bar = 45 μm.

The composition of the wall of the horns and the uterine body was similar, comprising three layers: the perimetrium, the myometrium and the endometrium (Figure 5 - B). The perimetrium is the outer serous layer consisting of mesothelium, a simple squamous epithelium that provides the lining of the organ, and a layer of loose connective tissue, where the fibers of its extracellular matrix are loosely arranged, arranged longitudinally along the length of the organ.

The myometrium is the intermediate layer composed of smooth muscle tissue, also known as non-striated muscle tissue or visceral muscle tissue, consisting of mononucleated and elongated cells, arranged in an inner circular layer and an outer longitudinal layer.

The endometrium is the inner layer of the uterus, also called the uterine lining. A simple columnar epithelium was presented, a single column-shaped layer of cells that extends within the endometrial stroma, forming tubular glands formed from the same simple columnar epithelium, as shown in figure 5 - C.

The endometrial stroma is composed of stromal cells, which form the supporting tissue of the epithelium, in addition to blood vessels in a loose connective tissue. There was a variation between the animals in relation to the height of the stroma, which is characteristic of the uterine dynamics during the female’s reproductive cycle.

Table 4 presents the mean values referring to uterine morphometry. It was possible to verify that the uterus has an average thickness of 725.26 μm, except for the thickness of the perimetrium, which it was not possible to measure in most of the analyzed specimens. The thickness of the endometrial wall had a wide variation in its values, ranging from 193 to 517 μm, which was reflected in the high standard deviation found. No external signs of menstruation were observed in the analyzed females and the histological analysis did not show any process of desquamation or necrosis in the endometrium. As for the thickness of the muscle layer, there was a variation between 151 and 280 μm, with an average of 212.27 μm. The inner circular muscle layer was thicker than the outer circular muscle layer, and both showed a lower standard deviation in terms of thickness, compared to the previously mentioned parameters.

**Table 4.**
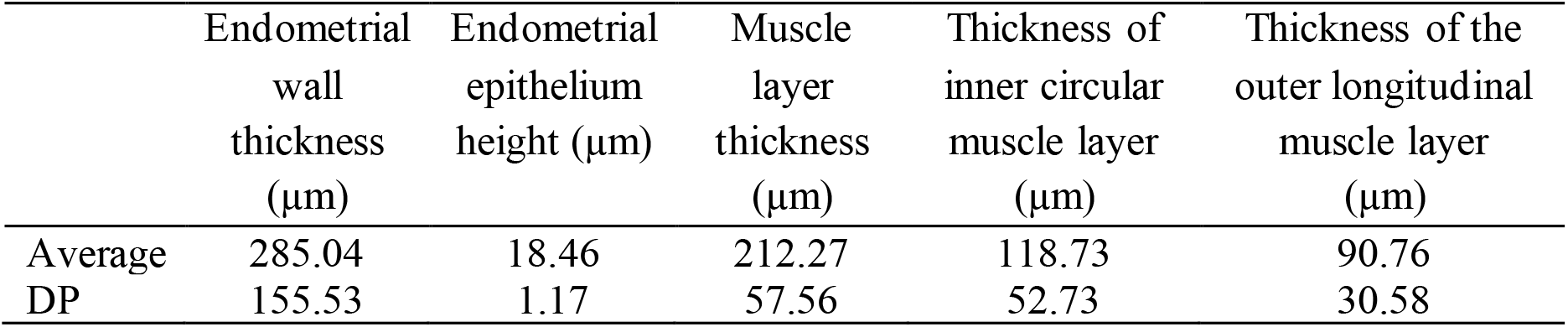
Components of the Uterus of the bat *Pteronotus gymnonotus*. Data expressed as mean ± standard deviation.

| | Endometrial<br>wall<br>thickness<br>($\mu\text{m}$) | Endometrial<br>epithelium<br>height ($\mu\text{m}$) | Muscle<br>layer<br>thickness<br>($\mu\text{m}$) | Thickness of<br>inner circular<br>muscle layer<br>($\mu\text{m}$) | Thickness of the<br>outer longitudinal<br>muscle layer<br>($\mu\text{m}$) |
| --- | --- | --- | --- | --- | --- |
| Average | 285.04 | 18.46 | 212.27 | 118.73 | 90.76 |
| DP | 155.53 | 1.17 | 57.56 | 52.73 | 30.58 |

## 5. DISCUSSION

This is the first study approaching morphological aspects of the female reproductive organs of the bat *P. gymnonotus*. Due to the difficulty in accessing its cave habitat, little is known about the reproductive biology of the species. In general, the morphology of the organs analyzed in this study was similar to the pattern described for other mammals.

### 5.1 OVARIAN MORPHOLOGY

The ovaries were constituted by the central region, formed by loose connective tissue, where blood and lymphatic vessels are found, in addition to interstitial cells and nerves. The peripheral part was wide and formed by connective tissue, where several follicles in various stages of maturation were observed. These follicles gradually migrate to the peripheral region of the ovary as they develop and increase in size. This pattern is also found in other bat species, such as *Pteropus giganteus*, *Artibeus fimbriatus* and *Artibeus obscurus* (Dorlikar et al., 2013).

However, some species of bats do not follow this same general pattern, as is the case of the phyllostomids *Artibeus planirostris*, *Sturnira lilium*, *Glossophaga soricina* and *Leptonycteris curasoae*, among others, which have an ovarian polarity in which a portion of the ovary is covered by epithelium and the follicles are restricted in this area until they are recruited at ovulation (Heideman and Powell, 1998; Antonio-Rubio et al., 2013).

This variation in the arrangement of ovarian follicles in different species of bats is related to the variety of reproductive patterns observed in the order Chiroptera, as well as to numerous specializations and peculiarities, such as asymmetry of Organs reproductive organs; characteristic where a single ovary is dominant (Wimsatt, 1979). This asymmetry pattern appears in several species of bats, the most common being the complete or partial dominance of the right ovary in species from the families *Molossidae*, *Rhinolophidae* and Mormoopidae; and dominance of the left ovary, in species of the *Emballonuridae* and *Megadermatidae families* (Pillai and Sastry, 2012). The ovaries of *P. gymnonotus*, in the specimens analyzed in this study, presented corpus luteum in almost 10% of all analyzed samples, which confirms the functionality of both ovaries (Vicente et al., 2006).

This ovarian asymmetry in bats has been interpreted by making a direct relationship with the number of offspring, which in general is only one per gestation; species that have more than one offspring per gestation are considered bilaterally symmetrical; that is, both ovaries are functional and ovulate at the same frequency (Pillai and Sastry, 2012). In this study, even though *P. gymnonotus* presented both functional ovaries, in which it presented mature follicles and corpus luteum in all analyzed ovaries, it was not possible to identify whether both ovulated at the same frequency or if they presented a certain dominance, so at the moment it was not possible to confirm whether the ovaries of the species can be considered bilaterally symmetrical.

The analysis of the folliculogenesis of *P. gymnonotus* demonstrated that the growth and development of the follicles followed the typical pattern of other mammals, as well as of other bats, such as *Miniopterus fraterculus* (Bernard, 1980), *Artibeus fimbriatus*, *Sturnira lilium* (Fazzolari-Corrêa, 1995) and *Pteropus giganteus* (Dorlikar et al., 2013). According to Adona (2012), the standard folliculogenesis process in mammals occurs in the ovarian cortex, where in the transition from mitosis to meiosis, oogonia transform into primary oocytes and primordial follicles, located on the periphery of the ovary. In most species, the granulosa cells of primordial follicles may originate from mesothelial cells and have two distinct formats: squamous and cubic, depending on their stage of development (Beguelini, 2020).

The follicle is responsible for producing the steroid hormones and proteins needed to maintain the ovarian cycle, secondary sexual characteristics and preparation for implantation in the uterus, and after ovulation, the corpus luteum supplies the hormones needed to maintain the pregnancy. The maintenance of this functional cycling pattern is controlled by pituitary gonadotropins (FSH and LH) and ovarian steroid hormones (estradiol and progesterone), which bind to specific receptors in the ovary (Findlay et al., 2009).

Primordial follicles remain dormant in the ovaries until they are recruited and grow to develop into a unilaminar primary follicle, when a single layer of flattened granulosa cells surround the oocyte. Its progression and the presence of another layer of granulosa cells, now cubic, causes the follicle to be called a primary multilaminar follicle, with the formation of the zona pellucida around the oocyte still occurring at this stage; which increases in size at that time (Adona, 2012).

Finally, through cell proliferation, the follicle begins to form an antrum due to the increase in cells and their cytodifferentiation, becoming a secondary follicle. Finally, at the end of its development and maturation, the follicle becomes mature, with granulosa cells forming the *corona radiata* and *cumulus oophorus regions*, as well as lining the antral cavity. It is at this time that the secondary oocyte emerges and can be released from the ovary at the time of ovulation (Erickson, 2002). The results of this study showed that females of *P. gymnonotus*, as well as other bat species (Fazzoli-Correa, 1995), are monovular, given that only one follicle has matured; while most of the rest progressed only to the multilaminar primary follicle stage.

As soon as the follicle reaches its maximum stage of maturation, that is, it becomes a mature follicle, its rupture occurs, where the oocyte is released. As a result, there is the formation of a single corpus luteum per follicle (Tomac et al., 2011). In the ovary of *P. gymnonotus* that presented the corpus luteum, large amounts of primordial follicles and smaller amounts of primary and secondary follicles were found; while in the ovary without corpus luteum, more secondary and mature follicles were observed, and a smaller number of primary follicles. Thus, the ovary without the corpus luteum was similar to other bat species, presenting some characteristics such as: relatively smaller size than the ovary with the corpus luteum and a significant decrease in its diameter. These similar characteristics demonstrate that there is a regression process undergone by this ovary, as a result of which ovary ovulated (Tomac et al., 2011).

The corpus luteum is a transient endocrine gland, responsible for the initial maintenance of pregnancy (Dorlikar et al., 2013). It secretes hormones, mainly progesterone, which will act to maintain the inner layer of the uterus, the endometrium, to receive and maintain the conceptus. The corpus luteum can remain until late pregnancy or early lactation and can occupy an area of up to 70% in ovarian section (Dorlikar et al., 2013). Thus, in the species *P. gymnonotus*, it presented a corpus luteum in its initial phase, where the granulosa cells invaded the antrum to start the process of transformation into luteal cells and start the production of hormones, mainly progesterone.

In the ovaries that presented corpus luteum, there was a greater thickening of the endometrial layer of this same animal, showing an increase in the number of glands, indicating the end of the ovulation phase and the beginning of the luteal phase of the ovarian cycle. However, ovaries that did not present a corpus luteum, but with follicles in growth and development, were associated with a thinner endometrium and a reduction of the glands, indicating a proliferative phase of the endometrial cycle; which was in line with observations made in ovaries with and without corpus luteum, along with variations in the thickness of the endometrium in the uterus (Bueno, 2019).

### 5.2 TUBE MORPHOLOGY

The histology of the uterine tubes of *P. gymnonotus* shows similarities to the pattern of mammals and as observed in bat species, with *Myotis nigricans*, family Vespertilionidae (Beguelini et al., 2020). Composed of three layers: the serous layer, which has the function of lining the organ, presenting connective tissue and covered by mesothelium; the thickest layer is the muscular layer, composed of smooth muscle cell, being represented by two layers, a longitudinal layer and another circular layer, and the innermost layer and with the function of performing the necessary contraction for the transport of the oocyte or zygote for uterus or uterine horns; the mucosa with non-ciliated (secretory) cells that secrete mucus into the uterine tubes and ciliated cells that move the mucus that covers their surface towards the uterus or uterine horns; and in which it was observed in all parts of the uterine tube, without significant differences. (Al-Saffar; Almayahi, 2019) (Abiaezute, Nwaogu, Okoye, 2017).

### 5.3 UTERINE MORPHOLOGY

Females of *P. gymnonotus* had a bicornuate uterus, as found in other species of bats, such as *Molossus molossus* (Martins et al., 2011). The uterus in mammals in general presents a diversified morphological variation, such as a double uterus, with completely separate horns and presenting two independent cervixes, the two horns being able to be interconnected by two cranial openings in the cervix; bipartite uterus, in which a discreet fusion in the inferior region of the horns is its main feature; bicornuate uterus, in which a good part of the uterine horns are fused; and simple uterus, containing a single cavity resulting from the complete fusion of the horns. The latter is found in most primates and also some species of bats in the *Phyllostomidae family*. However, bipartite and simple uteri have also been found in the order Chiroptera, and there seems to be a predominance of the bicornuate uterus (Vicente et al., 2006; Kardong, 2010; Martins et al., 2011).

In primates and macroscelids, the innermost layer of the uterus, the endometrium, undergoes morphological changes, increasing its thickness close to the period of ovulation and sloughing off if fertilization does not occur, leading to menstruation (Mayor et al., 2019). Among bat species, some show evidence of true menstruation, as in molossids and phyllostomids (Rasweiler and Bradwaik, 1996; Komar, et al., 2007). A common feature among species that have true menstruation is that the vast majority of them have a simple uterus (Rasweiler and Bradwaik, 2000). Histological analysis of the *P. gymnonotus uterus* did not show signs of endometrial desquamation, or even signs of bleeding from the vulva at the time of collection; although true menstruation has been linked to bats with a bicornuate uterus, such as *Molossus rufus* and *Rousettus leschenautii* (Catalini and Fedder, 2020). Thus, new studies are needed to allow understanding about the morphophysiology of the endometrial cycle in *P. gymnonotus*.

The results obtained showed that the endometrial cycle was intrinsically linked to the ovarian cycle. The uterus of *P. gymnonotus* underwent some changes that coincided with some phases of the ovarian cycle, with emphasis on the thickening of the endometrial layer during the end of the ovulation phase and the beginning of the luteal phase, indicating the secretory phase of the endometrial cycle, since in this phase, the endometrium obtained the highest values in terms of thickness, with a greater number of glands. However, in ovaries that did not have a corpus luteum, but showed growing and developing follicles, the endometrium was smaller and a reduction in the number of glands, indicating the proliferative phase.

## 6. FINAL CONSIDERATIONS

The joint analysis of the results showed similar reproductive characteristics in *P. gymnonotus in relation* to other bats and, mainly, in view of the general characteristics of mammals, despite not having a polarization of the ovaries and not actually presenting a bilateral asymmetry, with dominance of one ovary. As for the folliculogenesis process, *P. gymnonotus* presents the same basic pattern of mammals in general, in particular, of other bats. There was a relationship between the phases of the ovarian and endometrial cycles, highlighting the period of ovulation, in the transition from the proliferative phase to the secretory phase of the ovarian cycle, with the secretory phase of the endometrial cycle.

